# HOPS is required for non-canonical mTORC1 signaling by recruiting Rags and FLCN to lysosomal membranes

**DOI:** 10.64898/2026.05.14.724810

**Authors:** Reini E. N. van der Welle, J.A. van der Beek, P. Sanza, F.J.T. Zwartkruis, N. Liv, J. Klumperman

## Abstract

The mechanistic target of Rapamycin complex 1 (mTORC1) regulates cell growth and metabolism in response to nutrient availability. mTORC1 recruitment to lysosomes by the Rag dimer complex (RagA/B and RagC/D) is a crucial step in mTORC1 signaling. Substrates of the MiT/TFE transcription factor family, like TFEB and TFE3, directly interact with the Rag complex when activated by folliculin (FLCN). This selective recruitment and subsequent phosphorylation of MiT/TFE substrates leads to a non-canonical mTORC1 signaling pathway regulating lysosomal biogenesis and autophagy. Recently, we showed that compound heterozygous mutations in VPS41 specifically impair non-canonical mTORC1 signaling. VPS41, as part of the HOPS complex, is required for fusion of lysosomes with endosomes and autophagosomes. Here we addressed the mechanism by which VPS41/HOPS complex regulates mTORC1 activity. We show that multiple HOPS subunits interact with the Rag dimers as well as with FLCN. Depletion of HOPS subunits results in reduced lysosomal localization of both Rags and FLCN, and the nuclear translocation of TFE3/TFEB. The VPS41-Rag interactions required the RING domain of VPS41, but were independent of RHEB, Rag activity or presence of other HOPS components. We conclude that HOPS is necessary for the recruitment of crucial components of the non-canonical mTORC1 signaling pathway onto lysosomal membranes. These data further our molecular understanding of disease-causing VPS41/HOPS mutations and indicate a crucial role for HOPS in connecting lysosomal trafficking to lysosomal signaling.

## Introduction

Lysosomes are membrane-bounded organelles that contain hydrolytic enzymes necessary for the degradation and recycling of macromolecules(Klumperman & Raposo, 2014; Luzio et al., 2007; Saftig & Klumperman, 2009). Positioned at the crossroad of the endocytic and autophagic pathways, lysosomes receive information from the extra- and intracellular environment(Inpanathan & Botelho, 2019; Reggiori & Klumperman, 2016), thus functioning as a uniting platform for various nutritional clues. mTOR is a 298 kD serine/threonine protein kinase which, as part of the mTORC1 complex, controls cell metabolism and growth. By integrating information on the availability of a.o. growth factors, amino acids, cholesterol, oxygen and energy, mTORC1 regulates the balance between protein synthesis and degradation(Rabanal-Ruiz & Korolchuk, 2018; Sancak et al., 2010; Shin et al., 2022).

Activation of the canonical mTORC1 pathway requires binding to Ras Homolog Enriched in Brain (RHEB), which results in an activating conformational change. RHEB-dependent mTORC1 activation mainly occurs on the lysosomal membrane, to which mTORC1 is recruited by a complex of ‘Ras related small guanosine triphosphate (GTP)-binding proteins’ (Rags). RagA/B in a heterodimeric complex with RagC/D associates with lysosomes by binding to the lysosomal Ragulator complex. In concert with amino acid sensors such as the v-ATPase and SLC38A9, Ragulator acts as guanine exchange factor (GEF) for RagA/B activation. Amino acids produced by lysosomal degradation stimulate Ragulator-controlled activation of RagA/B, thereby linking amino acid availability to RagA/B activation(Brady et al., 2016; Fromm et al., 2020; Jewell et al., 2014; Sancak et al., 2008, 2010). Activated mTORC1 drives cell growth via phosphorylation of the ribosome-associated eukaryotic translation Initiation Factor 4E (eIF4E)-Binding Protein 1 (4E-BP1) and Ribosomal protein S6 kinase beta-1 (S6K1)(Holz et al., 2005; Wu et al., 2017). Active mTORC1 also inhibits macro-autophagy (hereafter referred to as autophagy) by phosphorylation/inhibition of Atg proteins (e.g. ULK 1/2)(Deleyto-Seldas & Efeyan, 2021) and members of the MiT/TFE family of transcription factors (e.g. TFE3 and TFEB). When mTORC1 is inactivate, non-phosphorylated TFE3/TFEB translocate to the nucleus, where they promote expression of the Coordinated Lysosomal Expression and Regulation (CLEAR) gene network, inducing autophagy and lysosome biogenesis(Napolitano & Ballabio, 2016; Palmieri et al., 2011; Sardiello et al., 2009; Settembre et al., 2012).

Whereas mTORC1 signaling has long been considered as an on/off pathway affecting all substrates equally, emerging evidence points to a non-canonical mTORC1 signaling pathway that specifically controls the activity of the MiT/TFE transcription factors. This pathway is independent of RHEB and requires the direct interaction of MiT/TFE transcription factors with the Rag heterodimers on the lysosomal membrane(Martina et al., 2012; Napolitano et al., 2020; Puertollano et al., 2018; Wada et al., 2016). To allow this interaction, both Rags must be active, i.e. RagA/B in the GTP-bound and RagC/D in the GDP-bound state. RagA/B-GTP recruits mTORC1 to the lysosomal membrane, whereas RagC/D-GDP recruits the MiT/TFE substrates(Napolitano et al., 2020, 2022). This mechanism explains why stimuli that modify RagA/B activity – e.g. amino acids, glucose, cholesterol – affect all mTORC1 substrates, whereas inactivation of RagC/D – e.g. by lysosomal damage or selective autophagy – only affects phosphorylation of MiT/TFE transcription factors(Bar-Peled et al., 2013; Napolitano et al., 2020). The selective mechanism for lysosomal recruitment of MiT/TFE substrates thus leads to a non-canonical mTORC1 signaling pathway specifically regulating lysosomal biogenesis and autophagy.

Whereas 4E-BP1 and S6K1 phosphorylation is dependent on RHEB and growth factors, phosphorylation of TFE3 and TFEB requires Folliculin (FLCN) and amino acids. In complex with Folliculin-interacting protein 1 or 2 (FNIP1/2), FLCN induces RagC/D GTP to GDP conversion (i.e. activation)(Lawrence et al., 2019; Meng & Ferguson, 2018; Petit et al., 2013). Mutations in FLCN, causing the Birt-Hogg-Dube (BHD) syndrome, hence prevent phosphorylation of MiT/TFE substrates, leading to their constitutive activation(Lawrence et al., 2019; Napolitano et al., 2020). Other currently known factors that selectively impact this non-canonical mTORC1 signaling pathway are lysosomal damage, conditions of selective autophagy (mitophagy, pathogens), mutations in the tuberous sclerosis complex genes TSC1, TSC2 (which inhibit RagC/D via an unknown mechanism), and mutations in the ‘Homotypic Fusion and vacuole Protein Sorting’ (HOPS) complex(Sanderson et al., 2021; Van der Welle et al., 2021).

The HOPS complex consists of 6 subunits (*VPS11, VPS16, VPS18, VPS33A, VPS39* and *VPS41*) and mediates the fusion of lysosomes with late endosomes and autophagosomes, a crucial step in the transfer of substrates to enzymatically active lysosomes (Beek et al., 2019; Caplan et al., 2001; Kant et al., 2015; Pols et al., 2013; Seals et al., 2000; Wurmser et al., 2000). In a recent study we showed that mutations or depletion of HOPS subunit *VPS41* cause a decrease in lysosomal association of mTORC1 and a constitutive nuclear localization of TFE3, while phosphorylation of 4E-BP1 and S6K1 was not affected(Sanderson et al., 2021; Van der Welle et al., 2021). Similar observations were made for *VPS11, VPS18* and *VPS39* knock-out cells and primary fibroblasts obtained from patients with compound heterozygous mutations in *VPS41*(Van der Welle et al., 2021). These data indicate a regulatory role of HOPS in the non-canonical pathway of mTORC1 signaling, but the underlying mechanism has remained unknown. In this study we address how HOPS specifically regulates mTORC1 activity towards TFE3/TFEB substrates. We show that the HOPS complex interacts with the Rag dimers as well as with FLCN and enables Rag recruitment and retention to the lysosomal membrane. Our study thus highlights a dual role for HOPS as tethering complex for endo-lysosomal membrane fusion and as a molecular determinant for lysosome-associated mTORC1 signalling.

## Results

### HOPS-dependent TFE3/TFEB phosphorylation is independent of RHEB

Previously, we addressed the effect of HOPS depletion on mTORC1 signaling in HeLa CRISPR/Cas9 cells knockout (KO) for the HOPS-specific components VPS41 or VPS39, or for VPS18, which is shared with the related CORVET complex(Beek et al., 2019; Kant et al., 2015). We showed by western blotting that in HeLa wildtype (WT) and HOPS KO cells cultured under basal, untreated conditions (i.e., in presence of serum), S6K and 4E-BP1 are phosphorylated. In HeLa WT and KO cells, phosphorylation of both mTORC1 substrates was decreased after 2 hours nutrient starvation and was restored after 15 minutes refeeding (restimulation) (Supplemental Fig 1A, B) and (Van der Welle et al., 2021). Nuclear localization of TFE3 in WT cells was only seen after nutrient starvation. By contrast, HeLa KO cells showed a constitutive nuclear localization of TFE3, irrespective of nutrient status, indicating that TFE3 phosphorylation depends on the presence of the investigated HOPS subunits (Supplemental Fig 1C, D) and (Van der Welle et al., 2021). Immunofluorescence of endogenous TFEB is difficult, but by western blot TFEB phosphorylation status is accompanied by an upward molecular weight shift(Roczniak-ferguson et al., 2012). In HeLa WT cells we found a clear shift of TFEB under fed conditions compared to starved conditions. By contrast, cells depleted of HOPS subunits never showed the upward shift, indicating that – like TFE3 – TFEB is not phosphorylated in HOPS KO cells (Supplemental Fig 1E). Moreover, we found an overall reduction in TFEB levels in the HOPS KO cell lines, indicating a negative feedback loop initiated by the constitutive activity of MiT/TFE transcription factors (Supplemental Fig 1E). Together these data show that phosphorylation of the canonical mTORC1 substrates S6K and 4E-BP1 is unaffected in HOPS KO cells, whereas phosphorylation of TFEB/TFE is blocked (Supplemental Fig 1) and (Van der Welle et al., 2021).

Previous studies have shown that overexpression of RHEB or siRNA-mediated silencing thereof significantly affects S6K1 and 4E-BP1 phosphorylation but has no effect on TFE3/TFEB phosphorylation(Martina et al., 2012; Puertollano et al., 2018; Wada et al., 2016). This predicts that RHEB over-expression will not be able to rescue TFE3/TFEB phosphorylation in HOPS depleted cells. To test this, we transiently transfected HeLa WT and VPS41^-/-^ cells with constitutively active RHEB (RHEB^N153T^). In both cell lines, expression of RHEB^N153T^ resulted in the constitutive phosphorylation of S6K1, even upon starvation (Fig 1A, B), nicely confirming the regulatory role of RHEB in phosphorylation of the canonical substrates. By contrast, over-expression of RHEB^N153T^ did not prevent nuclear localization of TFE3 upon starvation (Fig 1C), nor was able to rescue nuclear localization of TFE3 under basal conditions observed in VPS41^-/-^ cells. These data are in agreement with a RHEB-independent regulation of mTORC1 activity towards TFE3 (Napolitano et al., 2020) and show that the HOPS-dependent regulation of TFE3 phosphorylation is also independent of RHEB.

**Figure 1.**
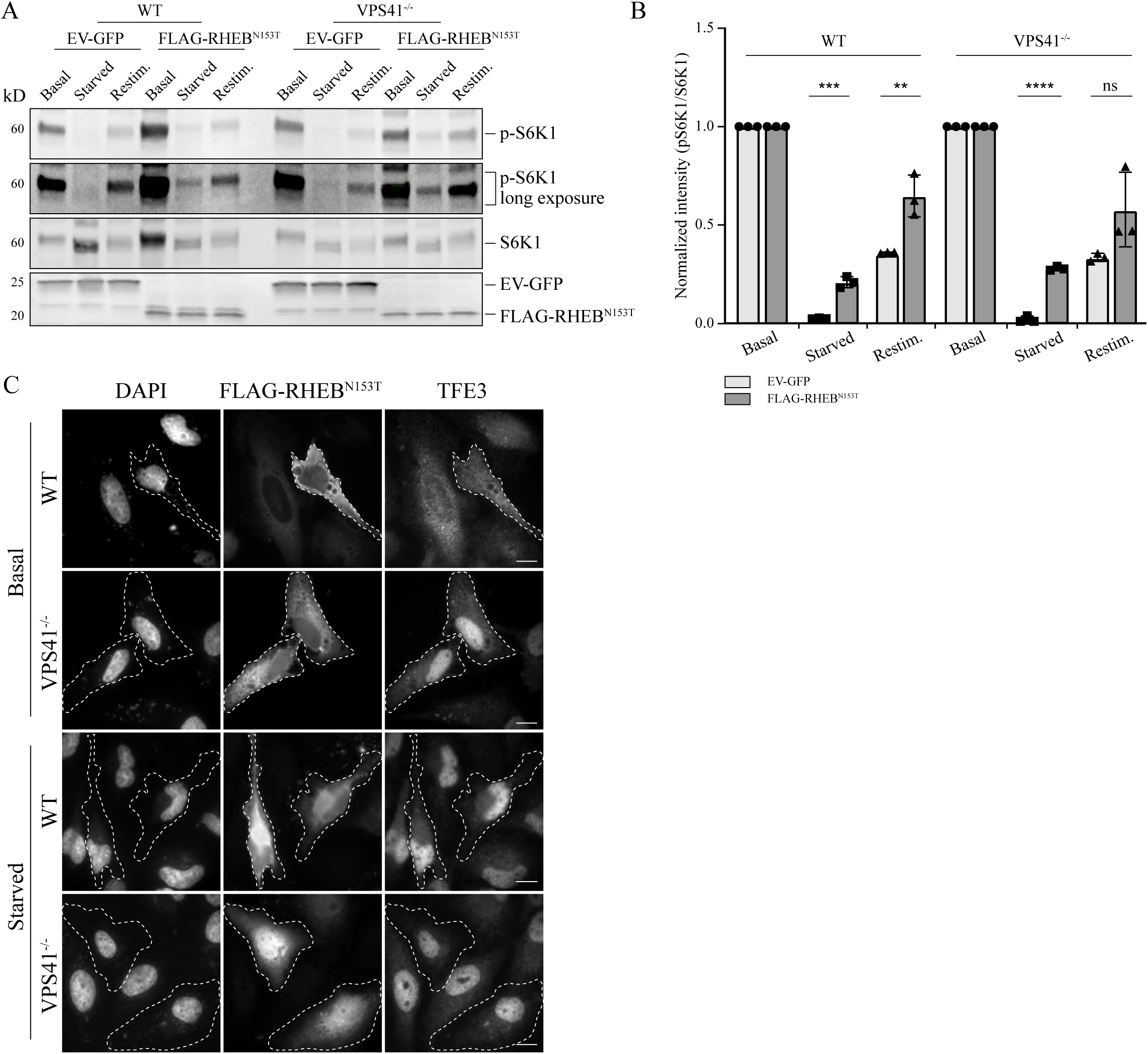
HOPS-dependent TFE3/TFEB phosphorylation is independent of RHEB. A. Western blot on the phosphorylation status of S6K1 in WT and VPS41^-/-^ cells overexpressing empty vector (EV-GFP) as negative control or constitutively active RHEB (FLAG-RHEB^N153T^). S6K1 phosphorylation levels were similar for both cell lines. The long exposure blots showed remaining phospho-S6K1 in starved conditions upon overexpression of RHEB^N153T^. B. Quantification of A, numbers represent mean ± SEM, **P < 0.01, ***P < 10-4, ****P < 10-5. One-way ANOVA with Bonferroni correction) (n=3). C. Immunofluorescence of endogenous TFE3 in starved WT or VPS41^-/-^ cells overexpressing constitutively active RHEB (FLAG-RHEB^N153T^). Overexpression of RHEB^N153T^ did not rescue nuclear localization of TFE3 (transfected cells visualized by white outline, n=2).

### The HOPS complex interacts with FLCN and is required for its lysosomal association

Phosphorylation of non-canonical substrates by mTORC1 is dependent on the FLCN:FNIP1/2 complex, the GAP required for activation of RagC/D (Bar-Peled et al., 2013; Castellano et al., 2017; Lawrence et al., 2019; Meng & Ferguson, 2018; Petit et al., 2013; Sancak et al., 2010). Under nutrient-poor, i.e., amino acid depleted, conditions, FLCN:FNIP1/2 binds the inactive RagA/B^GDP^ heterodimer (Bar-Peled et al., 2012; Lawrence et al., 2019; Tsun et al., 2013; Wada et al., 2016). Upon amino acid availability, FLCN stimulates GTP hydrolysis on RagC/D, generating the active RagA/B^GTP^-RAGC/D^GDP^ confirmation. In its active configuration (RagA/B^GTP^ - RagC/D^GDP^), the Rag complex recruits mTORC1 to lysosomes by interacting with the Raptor subunit of mTORC1(Sancak et al., 2008, 2010). FLCN-FNIP-dependent activation of the Rag dimers also allows binding of TFE3/TFEB, making this alternative, RHEB-independent, mechanism for mTORC1 activation a way to specifically regulate TFE3/TFEB recruitment and phosphorylation(Hong et al., 2010; Ramirez Reyes et al., 2021).

The phenotype of the HOPS subunit depleted cells strongly resembles that of cells lacking FLCN, i.e. non-phosphorylated and nuclear TFE3 in combination with normal regulation of S6K1 and 4E-BP1 (Supplemental Fig 1)(Fromm et al., 2020; Napolitano et al., 2020). This raises the possibility that HOPS on lysosomes is required for recruitment of FLCN and/or activation of its GAP activity towards RagC/D. To address this, we investigated the lysosomal localization of FLCN by immunofluorescence microscopy in HeLa WT and VPS41^-/-^ cells. Since FLCN only localizes to the lysosomal membrane upon starvation, cells were starved for 2 hours prior to fixation. Then cells were double-labeled for endogenous FLCN and the late endosome/lysosomal marker LAMP1 either in the presence (Fig 2A) or absence (Supplemental Fig 2) of Vacuolin (5µM, 1 hour)(Cerny et al., 2004), a PIKfyve inhibitor that causes lysosomal swelling. In our hands, and in previous studies, the use of Vacuolin resulted in a better FLCN visualization(Cerny et al., 2004; Petit et al., 2013). Of note, the FLCN antibody yields a strong nuclear signal. Although a small nuclear pool of FLCN cannot be ruled out, previous studies showed non-specificity of this nuclear signal(Meng & Ferguson, 2018). Therefore, the antibody is deemed suitable to assess lysosomal recruitment of endogenous FLCN. Analysis of the fluorescent data showed that depletion of VPS41 resulted in a strikingly reduced co-localization between LAMP1 and FLCN (Fig 2B), indicating a role for VPS41 in lysosomal recruitment of FLCN. We then performed co-immunoprecipitation studies on HeLa WT cells transfected with FLAG-FLCN and GFP-VPS18, VPS39-V5 or GFP-VPS41. Interestingly, all these HOPS subunits were able to co-immunoprecipitate FLCN (Fig 2C), indicating that multiple HOPS subunits can, directly or indirectly, interact with FLCN. Collectively these data show that the presence of HOPS subunits on lysosomes is required for lysosomal association of FLCN.

**Figure 2.**
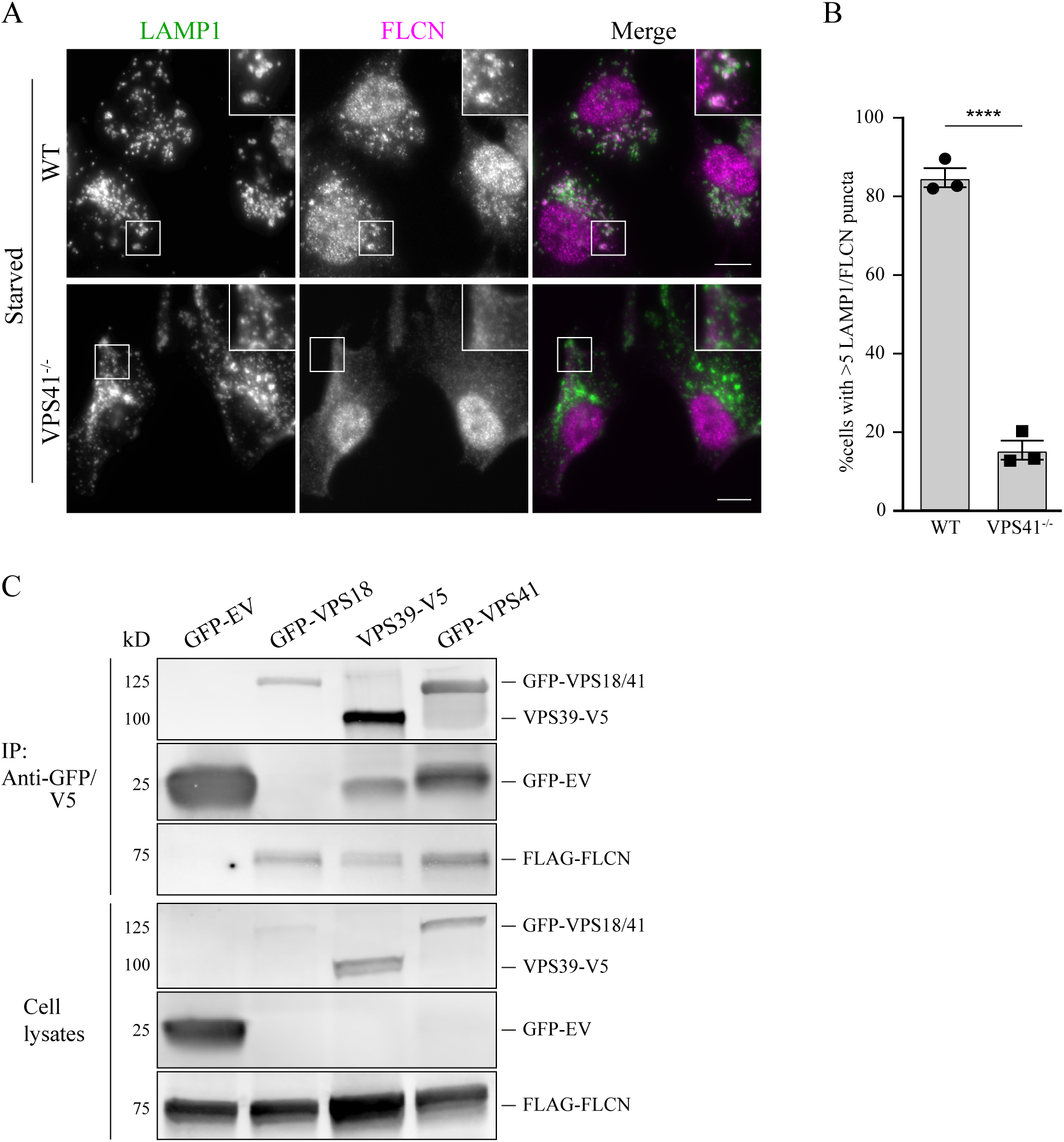
The HOPS complex interacts with FLCN and is required for its lysosomal localization. A. Immunofluorescence of endogenous FLCN and LAMP1 in starved, WT or VPS41^-/-^ cells treated with Vacuolin-1 (5µM, 1hour) to enlarge endo-lysosomal membranes (see Supplemental Fig 2 for FLCN localization in untreated cells). In VPS41^-/-^ cells, FLCN co-localization with LAMP1-positive compartments was significantly decreased. B. Quantification of A, showing percentage of cells with 5 or more FLCN spots that were also LAMP1 positive; mean ± SEM, ****P < 10-5. unpaired t-test). (Scale bar, 10 µm, n=3). C. Immunoprecipitation (IP) on cells co-expressing GFP-empty vector (GFP-EV), GFP-VPS18, V5-VPS39 or GFP-VPS41 and FLAG-FLCN. All three HOPS complex subunits interacted with FLCN.

### The HOPS complex is required for lysosomal recruitment of RagC

FLCN is recruited to lysosomal membranes by binding to the inactive RagA/B^GDP^ dimer of the heterodimeric Rag complex (Petit et al., 2013). The impaired lysosomal association of FLCN in HOPS KO cells could therefore indicate a lack of inactive Rag complex on lysosomes. To investigate lysosomal association of Rags in HOPS depleted cells, HeLa WT, VPS18^-/-^, VPS39^-/-^ and VPS41^-/-^ cells were starved for 0, 15, 30, 60, 90 or 120 minutes to initiate Rag recruitment. After fixation cells were double-labeled for immunofluorescence microscopy of RagC and LAMP1 (Fig 3A and Supplemental Fig 3A). RagC is the only Rag subunit that can be detected at endogenous levels, and since there are no indications that Rags dissociate into monomers, we considered this representative for the entire Rag complex(Sancak et al., 2008). In Hela WT cells cultured under basal conditions, we found very little co-localization between RagC and LAMP1 After 15 minutes starvation, however, co-localization strongly increased and after longer starvation, up to 90% of RagC positive puncta co-localized with LAMP1 (Fig 3A, B and Supplemental Fig 3A). Strikingly, in VPS18^-/-^, VPS39^-/-^ or VPS41^-/-^ cells, lysosomal recruitment of RagC was significantly reduced to a maximum of ± 68% co-localization after 120 minutes starvation (Fig 3A, B and Supplemental Fig 3A). As lysosomal localization of Rags is expected under basal conditions in WT cells, we verified these results using a different antibody to detect endogenous RagC in WT and VPS41^-/-^ cells under basal and starved conditions. Even though more co-localization between RagC and LAMP1 was detected in WT cells under basal conditions, the observed RagC – LAMP1 co-localization in VPS41^-/-^ cells was comparable between the two antibodies and significantly lower compared to WT cells (Supplemental Fig 3B).

**Figure 3.**
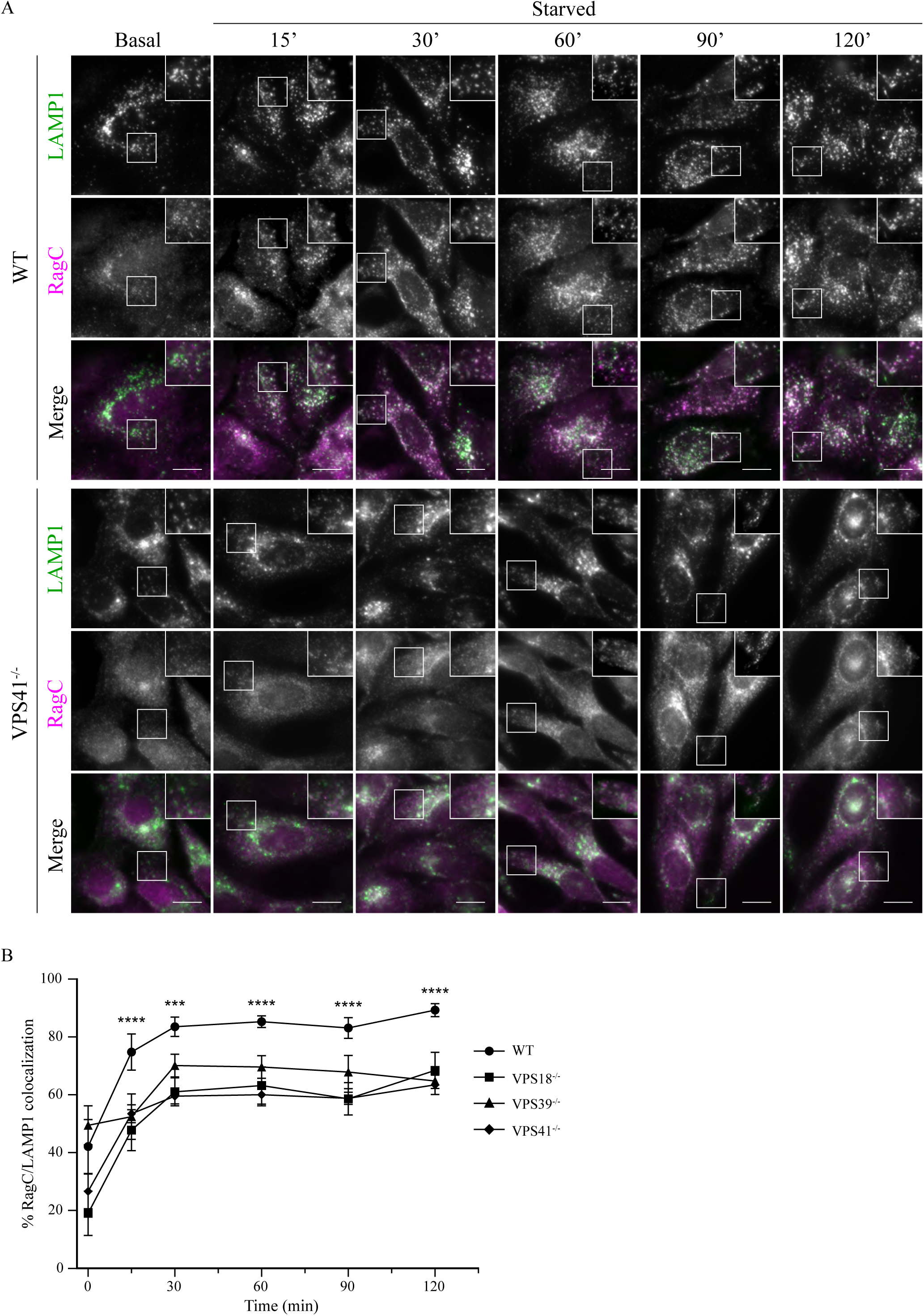
The HOPS complex is required for lysosomal recruitment of RagC. A. Immunofluorescence of endogenous RagC and LAMP1 in WT and VPS41^-/-^ cells under basal conditions and after different time points of starvation. In WT cells, RagC / LAMP1 colocalization markedly increased after 15 min or longer starvation. In VPS41^-/-^ cells, RagC remained cytosolic, showing little colocalization with LAMP1. Similar results were obtained for VPS18^-/-^ and VPS39^-/-^ cells (Supplemental Fig 3). (Scale bar and inserts, 15µm, n=2). B. Quantification of % RagC / LAMP1 co-localization in WT, VPS18^-/-^, VPS39^-/-^ or VPS41^-/-^ cells at different time points after starvation. In knockout cells this was significantly less at all time points (mean ± SEM, ***P < 10-4, ****P < 10-5. One-way ANOVA with Bonferroni correction) (n=2).

We conclude from these data that the HOPS complex is required for efficient lysosomal association of the Rag GTPases. Furthermore, the reduction in lysosomal association of Rags in HOPS depleted cells is a plausible explanation for the reduced lysosomal recruitment of FLCN (Fig 2A, B and Supplemental Fig 2).

### The HOPS complex is required for Rag-dependent lysosomal recruitment of TFE3

Besides FLCN and mTORC1, also TFE3/TFEB interact with the Rag complex present on lysosomal membranes. To establish if the reduced lysosomal localization of Rags in HOPS depleted cells (Fig 3A, B) explains the loss of lysosomal TFE3, we treated HeLa WT, VPS18^-/-^, VPS39^-/-^ and VPS41^-/-^ cells with Torin-1 and visualized lysosomal recruitment of TFE3 by immunofluorescence microscopy. Torin-1 suppresses mTORC1 kinase activity, inducing accumulation of non-phosphorylated TFE3 at the lysosomal membrane(Martina & Puertollano, 2013). Consequently, Torin-1 allows detection of lysosomal TFE3 not seen in non-treated cells. In untreated HeLa WT cells double-labeled for endogenous LAMP1 and TFE3, we found TFE3 diffused in the cytosol with no apparent enrichment on LAMP1-positive compartments (Fig 4A). By contrast, and in agreement with our previous observations, untreated VPS18^-/-^, VPS39^-/-^ and VPS41^-/-^ cells showed a strong nuclear localization of TFE3 (Fig 4A and Supplemental Fig 4A). After Torin-1 treatment, in HeLa WT cells co-localization between LAMP1 and TFE3 significantly increased, indicating that a pool of non-phosphorylated TFE3 remained associated with lysosomes (Fig 4A). Concomitant with mTORC1 inhibition, Torin-1 treatment also resulted in a bit of nuclear staining of TFE3. Intriguingly, in the HOPS depleted cell lines treatment with Torin-1 did not induce any co-localization between TFE3 and LAMP1 (Fig 4A and Supplemental Fig 4A). These data indicate that cells depleted of HOPS subunits entirely fail to recruit TFE3 to lysosomes for phosphorylation by mTORC1.

**Figure 4.**
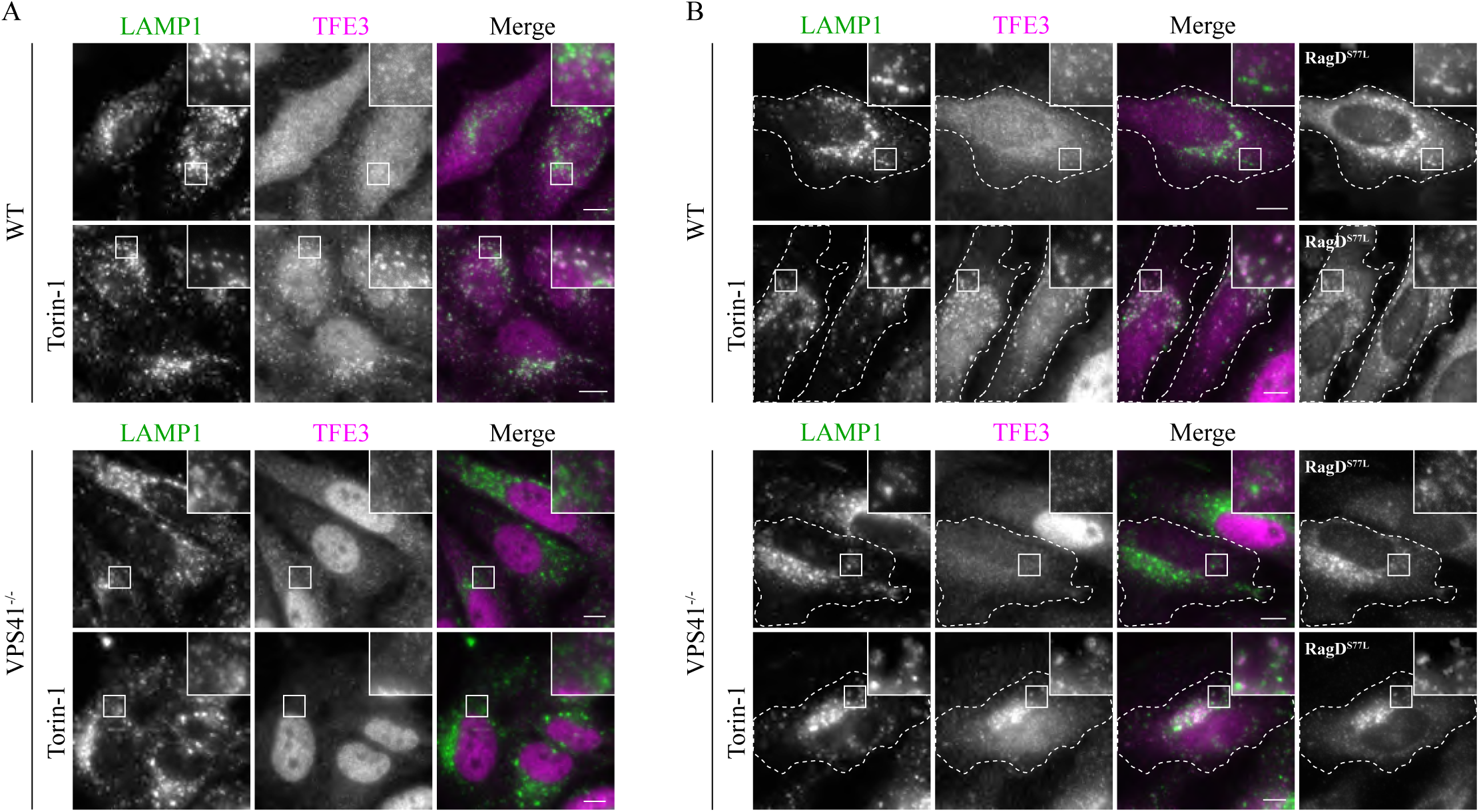
Lysosomal TFE3 localization is reduced in VPS41^-/-^ cells due to absence of active Rag GTPases. A. Immunofluorescence of endogenous TFE3 and LAMP1 in WT and VPS41^-/-^ cells with or without 250nM Torin-1 treatment for 1 hour. In WT cells, Torin treatment increased TFE3/ LAMP1co-localization, whereas in VPS41^-/-^ cells, TFE3 remained restricted to the nucleus. Similar results were obtained for VPS18^-/-^ and VPS39^-/-^cells (Supplemental Fig 4A). (Scale bar and insert, 10µm. n=3). B. Immunofluorescence of endogenous TFE3 and LAMP1 in WT and VPS41^-/-^ cells in the presence or absence of 250nM Torin-1 and upon transient transfection of constitutively active RagD (RagD^S77L^). In WT cells Torin-1 treatment induced co-localization of TFE3 with LAMP1 and RagD. Overexpression of RagD^S77L^ in VPS41^-/-^ cells rescued the nuclear localization of TFE3 and showed TFE3/LAMP1 co-localization on RagD positive compartments. Similar results were obtained for VPS18^-/-^ and VPS39^-/-^ cells (Supplemental Fig 4B) (transfected cells visualized by white outline). (Scale bar and insert, 10µm. n=3).

To further investigate the lack of Rag-lysosome association in HOPS KO cells, we transfected Hela WT, VPS18^-/-^, VPS39^-/-^ and VPS41^-/-^ cells with constitutively active RagD (HA-RagD^S77L^), which should rescue the nuclear localization of TFE3 in HOPS KO cells. Indeed, expression of HA-RagD^S77L^ completely abolished the constitutive nuclear localization of TFE3 in all HOPS KO cell lines and strikingly increased co-localization between LAMP1 and TFE3 upon Torin-1 treatment (Fig 4B and Supplemental Fig 4B). Even upon nutrient starvation, TFE3 was not detected in the nuclei of HA-RagD^S77L^ expressing HOPS KO cells. Upon overexpression of HA-RagD^WT^, TFE3 remained in the nucleus regardless of nutrient availability in VPS41^-/-^ cells. This indicates that it is not the expression level but rather the activity of RagD that is required for TFE3 retention in the cytosol (Supplemental Fig 4C). Thus, over-expression of RagD^S77L^ rescued the phosphorylation defect and therefore constitutive activation of TFE3 in HOPS depleted cells. Together these data showed that lysosomal dissociation of TFE3 caused by HOPS deficiency is rescued by introducing active RagD.

Collectively, our data show a major impact of HOPS depletion on lysosomal association and activation of Rags, being a crucial initial step in the non-canonical mTORC1 signaling pathway.

### Patient-bound VPS41^R662*^ fails to recruit Rag heterodimers to lysosomes

Recently, we described 3 patients with compound heterozygous variants in the *VPS41* gene. These included a missense variant in the WD40 domain (VPS41^S285P^) and a nonsense variant in the Clathrin Heavy Chain Repeat (CHCR) domain (VPS41^R662*^)(Van der Welle et al., 2021). The latter mutation results in a truncated protein lacking the RING domain^29^, which is required to bind the other HOPS subunits. Notably, VPS41^R662*^ is still recruited to lysosomal membranes. VPS41^S285P^ does bind other HOPS subunits, but both VPS41^R662*^ and VPS41^S285P^ cause a delay in delivery of endocytosed cargo to lysosomes, indicating a defect in HOPS-dependent endosome – lysosome fusion. Importantly, patient-derived fibroblasts display a similar constitutive nuclear localization of TFE3 and reduced lysosomal localization of mTORC1 as seen in HOPS KO cells(Van der Welle et al., 2021).

To determine the effect of patient-bound VPS41 variants on lysosomal association of the Rag dimers, we generated HeLa VPS41^-/-^ cells stably expressing VPS41^WT^-GFP, VPS41^S285P^-GFP or VPS41^R662*^-GFP or H2B-mNeon as negative control (hereafter referred to as VPS41^-/-^-mNeon) (Supplemental Fig 5A). To study lysosomal recruitment of Rags, cells were starved for 2 hours and prepared for double-immunofluorescent labeling of endogenous RagC and LAMP1. As expected, VPS41^-/-^ and VPS41^-/-^-mNeon cells showed very little overlap between RagC and LAMP1, whereas VPS41^WT^-GFP cells displayed a similar level of RagC-LAMP1 co-localization as HeLa WT cells expressing VPS41^WT^-GFP (Fig 5A, B). Interestingly, introducing the VPS41^S285P^ variant rescued RagC/LAMP1 co-localization, whereas VPS41^R662*^ did not (Fig 5A, B). Concomitantly, stable expression of VPS41^WT^ or VPS41^S285P^ rescued the constitutive nuclear localization of TFE3 (Fig 5C, D), whereas TFE3 remained nuclear in both VPS41^-/--^- mNeon and VPS41^R662*^ expressing cells (Fig 5C, D). These data show that patient-bound VPS41^R662*^ is unable to recruit Rags to lysosomes and indicate the VPS41 C-terminal/RING domain as crucial determinator for Rag recruitment.

**Figure 5.**
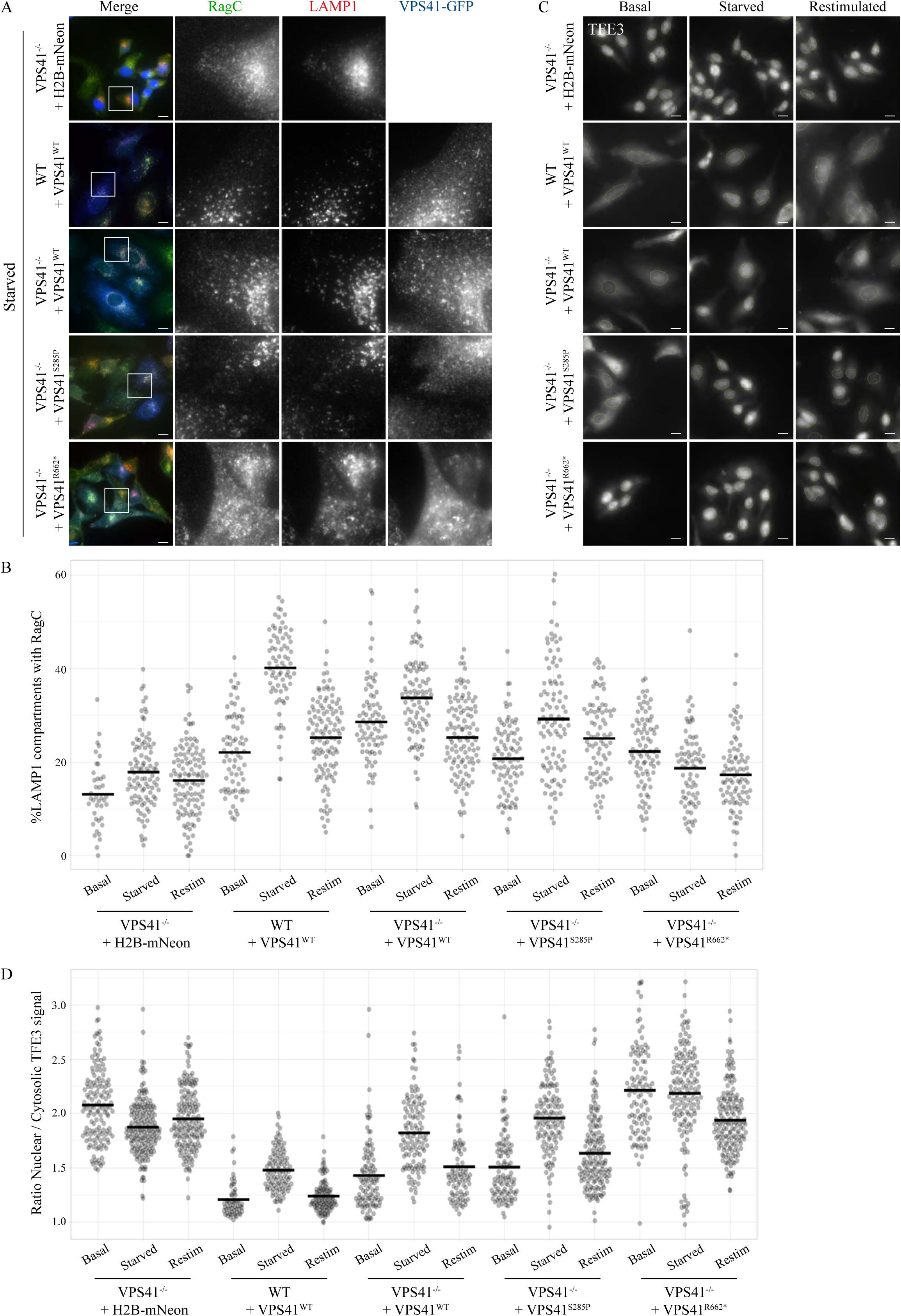
VPS41^R662*^ impairs lysosomal recruitment of RagC. A. Immunofluorescence labeling of endogenous TFE3, LAMP1 and RagC in starved VPS41^-/-^ cells stably expressing mNeon (negative control), VPS41^WT^-GFP, VPS41^S285P^-GFP or VPS41^R662*^-GFP. VPS41^-/-^-mNeon cells showed no colocalization between RagC and LAMP1. Stable expression of VPS41^WT^-GFP or VPS41^S285P^ - GFP rescued RagC recruitment to LAMP1-positive compartments, whereas stable expression of VPS41^R662*^ did not. (Scale bar, 10µm, Insert, 25µm. n= 2). B. Quantification of the RagC/LAMP1 co-localization upon stable expression of H2B-mNeon (negative control), VPS41^WT^-GFP, VPS41^S285P^-GFP or VPS41^R662*^-GFP. Expression of VPS41^WT^ and VPS41^S285P^ resulted in increased RagC/LAMP1 colocalization. C. Immunofluorescence of endogenous TFE3 in WT cells stably expressing VPS41^WT^ or VPS41^-/-^ cells stably expressing mNeon (negative control), VPS41^WT^-GFP, VPS41^S285P^-GFP or VPS41^R662*^-GFP under basal, starved (2 hours) or restimulated (15min) conditions. Knockout of VPS41 (VPS41^-/-^-mNeon) resulted in constitutive nuclear localization of TFE3, regardless of nutrient availability, whereas WT cells showed cytosolic retention of TFE3 in basal and restimulated conditions. Cells stably expressing VPS41^WT^ or VPS41^S285P^ showed a (partial) rescue of the nuclear TFE3 phenotype, resulting in cytosolic retention under basal and restimulated conditions. Expression of VPS41^R662*^ did not rescue nuclear localization of TFE3. D. Quantification of C. Numbers represent the ratio of cytosolic over nuclear TFE3, showing the rescue of TFE3 localization in VPS41^-/-^ cells stably expressing VPS41^WT^ or VPS41^S285P^ (n> 100 cells per condition).

### VPS41-Rag interactions require the VPS41-RING domain but not HOPS

To further investigate the role of the VPS41-RING domain in Rag interactions, we co-transfected VPS41^-/-^ cells stably expressing VPS41^WT^-GFP or VPS41^R662*^-GFP with active (RagA^GTP^/RagC^GDP^ - RagB^GTP^/RagD^GDP^) or inactive (RagA^GDP^/RagC^GTP^ - RagB^GDP^/RagD^GTP^) heterodimers. Subsequent co-immunoprecipitations showed interaction between VPS41^WT^ with both the active and inactive Rag dimers (Fig 6A and Supplemental Fig 5B), whereas VPS41^R662*^ failed to co-immunoprecipate the Rag heterodimer (Fig 6A and Supplemental Fig 5B). Interestingly, co-immunoprecipitations with FLCN showed that VPS41^WT^ and VPS41^R662*^ bound FLCN to a similar extent (Fig 6B). These data showed that the VPS41 RING domain is important for binding to the Rags, but not FLCN.

**Figure 6.**
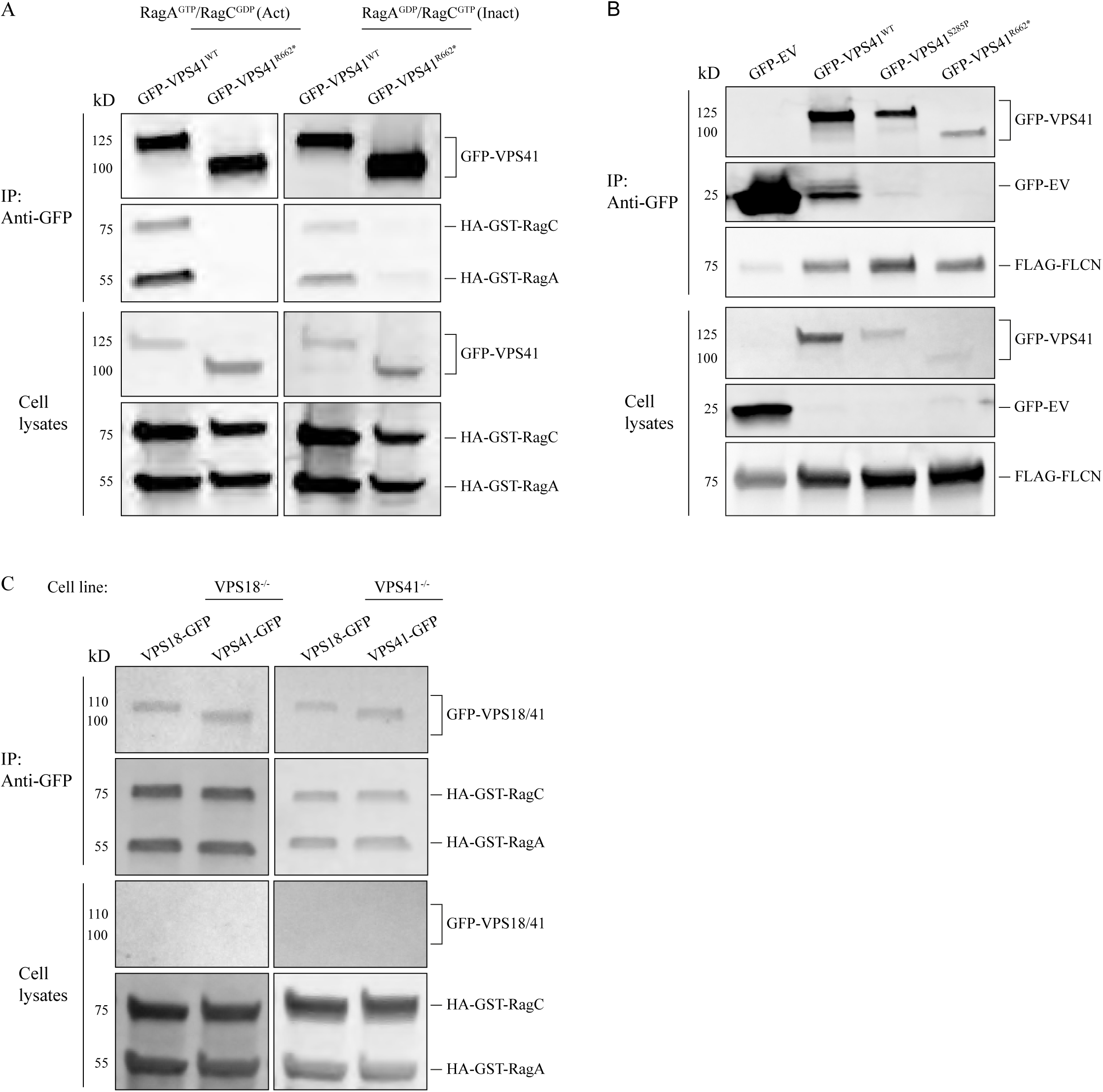
VPS41^R662*^ does not interact with Rags. A. Co-immunoprecipitation of cells stably expressing VPS41^WT^-GFP or VPS41^R662*^-GFP and transiently transfected with either RagA^GTP^/RagC^GDP^ (active conformation) or RagA^GDP^/RagC^GTP^ (inactive conformation). VPS41^WT^ interacts with both the active and inactive Rag heterodimers. The VPS41^R662*^ mutant, which lacks the RING domain, showed severely reduced affinity for the RagA/RagC heterodimer, regardless of conformation (n=3). B. Co-immunoprecipitation (IP) on cells co-expressing GFP-empty vector (GFP-EV), GFP-VPS41^WT^, GFP-VPS41^S285P^, GFP-VPS41^R662*^ and FLAG-FLCN. VPS41^WT^ as well as both mutants interact with FLCN (n=3). C. Co-immunoprecipitation (IP) on VPS18^-/-^ or VPS41^-/-^ cells co-expressing VPS18-GFP, VPS41-GFP and the RagA/RagC dimer. Both VPS18 and VPS41 interact with the RagA/RagC dimer in absence of the other HOPS subunit (n=3).

To establish if VPS41 – Rag interactions require binding of VPS41 to other HOPS subunits (via the RING domain), we next performed VPS41-Rag co-immunoprecipitation experiments in VPS18^-/-^ cells (Fig 6C). This yielded similar data as in VPS18-expressing cells (Fig 6A), indicating that the VPS41-RagA/C interaction does not require other HOPS subunits or formation of the HOPS complex (Fig 6C). Finally, to establish if other HOPS subunits also could bind Rags, VPS41^-/-^ cells were co-transfected with VPS18-GFP and RagA/C. Subsequent co-immunoprecipitations clearly showed an interaction between VPS18-GFP and the Rag dimers (Fig 6C), indicating that this interaction is not specific for VPS41.

Together these observations showed that the RING-domain of VPS41 is indispensable for interaction between VPS41 and FLCN but is required for interactions with the Rag dimers and essential for recruitment of Rags to lysosomes and subsequent phosphorylation of TFE3. In addition, VPS18 also interacted with Rags, indicating that multiple HOPS subunits may co-operate in the recruitment of Rags to lysosomes. How the HOPS complex exactly interacts with the Rag dimers on the lysosomal membrane, and how VPS41, VPS18, or possibly their RING domains are involved remains to be elucidated.

## Discussion

Recently we found that disease-causing mutations in VPS41 or depletion of HOPS components causes dissociation of mTORC1 from endo-lysosomal membranes and a constitutive nuclear localization of TFE3, while phosphorylation of 4E-BP1 and S6K1 was not affected(Sanderson et al., 2021; Van der Welle et al., 2021). This unexpectedly indicated a role for HOPS in the non-canonical mTORC1 signaling pathway(Napolitano et al., 2022). In this paper we investigated the mechanisms that underlie HOPS-dependent regulation of mTORC1 signaling. Our studies reveal that the presence of HOPS subunits on lysosomes is required to safeguard the first steps in non-canonical mTORC1 signaling, i.e. lysosomal association of Rags and FLCN, the GAP for activating RagC/D. We found that VPS41 interacted with the Rag heterodimer via its RING domain - which is absent from the previously identified, disease-causing VPS41^R662*^ mutant - and that this interaction is required for efficient recruitment of Rags to endo-lysosomes^29^. Binding of VPS41 to Rags did not require the presence of other HOPS subunits, but we did find that Rags also bind to VPS18. This suggests that a cooperative action of multiple HOPS subunits could be required for efficient Rag/FLCN recruitment to lysosomes. Altogether these findings reveal a dual role for HOPS at endo-lysosomal membranes; as tethering complex required for membrane fusion and as molecular interaction platform for mTORC1 signaling.

Depletion of the HOPS-core subunit VPS18 or the HOPS-specific components VPS39 or VPS41 all caused a significant reduction in co-localization of endogenous RagC and LAMP1 in nutrient depleted cells. In VPS41^-/-^ cells this phenotype, as well as the nuclear localization of TFE3, was rescued by reintroducing VPS41^WT^ or the patient-associated VPS41^S285P^ variant, but not by patient variant VPS41^R662*^, which lacks the RING domain. In agreement, VPS41^WT^ co-immunoprecipitated the RagA/RagC dimer, while this interaction was completely abolished in the VPS41^R662*^ variant. Overall, these data indicate that direct or indirect binding of the VPS41-RING domain to the Rag heterodimers mediates efficient lysosomal localization of the Rag complex. VPS41-Rag interactions did not depend on other HOPS subunits or assembly of the HOPS complex, since they were also observed in VPS18^-/-^cells. Furthermore, we found that also VPS18 binds Rags. Since both VPS18 and VPS41 bear a RING-domain, it is likely that the VPS18-RING domain is required for VPS18-Rag interactions. A third subunit of the HOPS complex, VPS11, also contains a RING domain. This raises the scenario that multiple HOPS subunits can bind Rags and co-operate to recruit the Rag complex to endo-lysosomal membranes.

Although interaction between VPS18 or VPS41 and Rags was observed in respectively VPS41^-/-^ or VPS18^-/-^ cells, the RagC and LAMP1 co-localization was significantly reduced in VPS18^-/-^ cells and VPS41^-/-^ cells. This indicates that interaction of the HOPS subunits outside the context of the tethering complex is not sufficient for endo-lysosomal Rag recruitment.

Interestingly, both WT and HOPS-subunit depleted cells reached a plateau in RagC-LAMP1 co-localization after 30 minutes of starvation (albeit at a lower level in the HOPS depleted cells), indicating that recruitment of Rags is not completely abolished, but more likely that dissociation from the lysosomal membrane is increased. Whether this is indicative of a change in kinetics, i.e., increased off rate in HOPS depleted cells remains to be elucidated. In addition, even though the independent HOPS subunits interact with the Rags, depletion of a single HOPS subunit is sufficient to impair efficient Rag retention/recruitment, demonstrating that a fully formed HOPS complex is required to establish a stable Rag complex on the lysosomal membrane.

Interestingly, like the Rags, the Rab-interacting Lysosomal protein (RILP) also interacts with the C-terminal region of VPS41(Lin et al., 2014). Therefore, interaction between RILP and VPS41^R662*^ will also be lost. RILP controls the transport and thus cellular distribution of endo-lysosomes by recruitment of dynein-dynactin motor complexes, preventing peripheral clustering of these compartments(Jordens et al., 2001; Kant et al., 2013). Additionally, FLCN interacts directly with RILP, thereby also playing a role in the perinuclear localization of endo-lysosomes(Starling et al., 2016). Since HOPS subunits interacted with FLCN, RILP could play a role in establishing the HOPS/Rag complex on the lysosomal membrane, where it also meets FLCN to regulate lysosomal positioning in response to nutrient status. Lysosomal positioning is previously linked to mTORC1 signaling(Korolchuk et al., 2011) and intriguingly, we have observed peripheral clustering of lysosomes in VPS41^-/-^ cells, indicative of impaired endo-lysosomal spatial distribution regulated by FLCN and RILP (data not shown).

The Rag complex is central in the non-canonical mTORC1 pathway by direct interactions with mTORC1 and members of the MiT/TFE transcription factor family. We show that in the absence of HOPS, Rags are not recruited or retained at endo-lysosomal membranes and propose that as a consequence, FLCN, mTORC1 and TFE3/TFEB also do not associate with endo-lysosomes. In accordance, we found that over-expression of constitutively active RagD rescued the phosphorylation and localization phenotype of TFE3 in VPS41^-/-^, VPS18^-/-^ and VPS39^-/-^ cells. Of note, VPS41^WT^ showed a strong affinity for both the active and inactive form of the Rag dimer. Since VPS41 binds both the Rags and FLCN, this might indicate a role for VPS41/HOPS in stabilization of the Rag/FLCN complex. In case both HOPS and the Rag complex are present, multiple direct or indirect interactions between HOPS subunits and Rags as well as with FLCN may help to strengthen or regulate their lysosomal association, allowing assembly of the Rag-mTORC1-MiT/TFE complex. In contrast to the Rags, VPS41-FLCN interaction did not require the VPS41-RING domain, indicating a role for the N-terminal domain of VPS41. The sequence of these interactions and if the interaction of HOPS with FLCN is Rag dependent or vice versa remains to be investigated.

Previously, we reported that patients bearing compound heterozygous mutations VPS41^S285P^ and VPS41^R662*^ suffer from a severe neurodegenerative disorder. Like HOPS KO cells, patient-derived fibroblasts show a reduced lysosomal association of mTORC1 and a constitutive nuclear localization of TFE3. Here we showed that stable expression of VPS41^S285P^ in VPS41^-/-^ cells increased lysosomal co-localization of RagC – LAMP1 and partially rescued the TFE3 nuclear localization. Since full VPS41 depletion is embryonically lethal(Aoyama et al., 2012), the observed residual functionality of the VPS41^S285P^ mutant is probably crucial for early survival of the VPS41^S285P^ / VPS41^R662*^patients. The partial rescue is in agreement with a recent paper describing that overexpressing of the VPS41^S285P^ mutation enables cytoplasmic localization of TFE3(Sanderson et al., 2021), but in seeming contrast to our own previous finding that transient expression of VPS41^S285P^ in VPS41^-/-^ cells did not rescue the TFE3 phenotype(Van der Welle et al., 2021). Most likely, these contradictory findings are explained by the difference in reintroducing the VPS41^S285P^ mutant. Transient expression can cause lysosomal stress, a known inducer for suppression of the non-canonical mTORC1 pathway and subsequent TFE3/TFEB activation(Martina et al., 2014). Lysosomal stress is avoided by making stable expressing cells, as performed in this study. This shows that transient expression experiments should be interpreted with caution when studying lysosomal stress.

As recently postulated by Napolitano et al., RHEB is required for S6K1 and 4E-BP1 phosphorylation, but dispensable for the regulation of TFE3/TFEB activity(Napolitano et al., 2020). Furthermore, regulation of S6K1/4E-BP1 phosphorylation is more sensitive to growth factors and TFE3/TFEB phosphorylation to amino acids availability, providing the means for independent regulation of canonical and non-canonical mTORC1 signaling. In agreement herewith, we showed that expression of constitutively active RHEB^N153T^ resulted in constitutive phosphorylation of S6K1 and 4E-BP1, but could not overcome the nuclear localization phenotype of TFE3 in HOPS depleted cells. RHEB induces a conformational change on mTORC1 necessary to obtain activity towards S6K1 and 4E-BP1(Yang et al., 2017), but not for TFE3/TFEB phosphorylation. A brief association of mTORC1 with the lysosomal membrane is probably sufficient for RHEB-dependent activation. Upon dissociation to the cytosol, RHEB-activated mTORC1 remains active and can phosphorylate substrates like S6K1 and 4E-BP1 by binding their TOS motif(Nojima et al., 2003; Schalm et al., 2003; Schalm & Blenis, 2002). By contrast, phosphorylation of non-canonical mTORC1 substrates is independent of RHEB activation and requires the presence of both mTORC1 and its substrates on the endo-lysosomal membrane(Jansen et al., 2022; Napolitano et al., 2020, 2022). Since both mTORC1 and TFEB/TFE3 interact with active Rags, their association with Rags present on the endo-lysosomal membranes is required to bring them together to allow phosphorylation of the TFEB/TFE3 transcription factors(Martina & Puertollano, 2013). Our data indicate that the presence of the HOPS complex results in longer association times of Rags on endo-lysosomes, thereby increasing the probability of TFE3/TFEB phosphorylation.

HOPS is required for endo-lysosomal fusion, which is necessary for the degradation of endocytic or autophagocytosed cargo(Beek et al., 2019; Jiang et al., 2014; Solinger & Spang, 2013) and the subsequent supply of amino acids that are crucial to locally activate the Rag complex via FLCN-FNIP(Gollwitzer et al., 2022). To enable a rapid response upon nutrient availability, FLCN binds to inactive Rags (i.e., under nutrient deprived conditions) on the endo-lysosomal membrane, thereby preparing the required machinery necessary for mTORC1 activation and TFE3/TFEB phosphorylation(Fromm et al., 2020; Jansen et al., 2022; Meng & Ferguson, 2018). The role for the HOPS complex proposed in this study ensures the formation of the Rag/FLCN complex on HOPS-positive endo-lysosomes. In addition, by mediating fusion of endosomes with active lysosomes, HOPS ensures the provision of amino acids necessary for FLCN to activate the Rags. Together these findings indicate a dual role for the HOPS complex in the activation of Rags; as a molecular platform to anchor the Rag/FLCN complex and as trafficking complex mediating the crucial fusion step of endosomes with active lysosomes, resulting in the supply of amino acids that leads to FLCN-dependent Rag activation.

Summarizing, our data demonstrate a major impact of HOPS depletion on the lysosomal association and activation of Rags. Since this is a crucial step in the lysosomal recruitment of FLCN and TFE3/TFEB, HOPS depletion consequently results in the selective inhibition of the non-canonical mTORC1 signaling pathway. Moreover, as an interactor with Rags and as a crucial factor for endo-lysosomal fusion, HOPS positions Rags precisely at the site where amino acids are generated (the active lysosome), enabling a rapid response upon amino acid availability and directly linking membrane trafficking to molecular signaling. Since an increasing number of pathological phenotypes are associated with dysregulation of mTORC1 and the MiT/TFE family of transcriptions factors, targeting the HOPS complex could be a promising therapeutic strategy to selectively modulate mTORC1 activity without interfering with the canonical signaling pathways.

## Methods and Material

### Antibodies and reagents

Antibodies and reagents used in this study and their specific dilutions or used concentrations are specified in Table 1.

**Table 1.** Antibodies and reagents used.

| Primary Antibody | Use | Dilution | Company | Catalog Number |
| --- | --- | --- | --- | --- |
| R α 4E-BP | WB | 1:1000 | Cell Signaling | 9644 |
| R α phospho-4E-BP1 (Ser65) | WB | 1:1000 | Cell Signaling | 9451 |
| R α p70 S6 | WB | 1:1000 | Cell Signaling | 9202 |
| R α phospho-p70 S6 (Thr389) | WB | 1:1000 | Cell Signaling | 9205 |
| R α TFEB | WB | 1:1000 | Cell Signaling | 4240 |
| M α VPS41 | WB | 1:500 | Santa Cruz | SC-377271 |
| M α Actin | WB | 1:2000 | MP Biomedicals | 69100 |
| R α GFP | WB | 1:1000 | Invitrogen | A6455 |
| M α GFP | WB | 1:2000 | Roche | 11814460001 |
| R α FLAG | WB | 1:2000 | Sigma | F7429 |
| G α HA | WB | 1:2000 | Genscript | A00168-100 |
|  | IF | 1:200 |  |  |
| R α V5 | WB | 1:1000 | Sigma | V8137 |
|  | IF | 1:400 |  |  |
| M α V5 | WB | 1:1000 | Invitrogen | R960-25 |
|  | IF | 1:200 |  |  |
| R α TFE3 | IF | 1:250 | Cell Signaling | 14779 |
| M $\alpha$ LAMP-1 CD107a | IF | 1:250 | BD Pharmingen | 555798 |
| R $\alpha$ FLCN | IF | 1:200 | Cell Signaling | 3697 |
| R $\alpha$ RagC (D8H5) | IF | 1:200 | Cell Signaling | 9480 |
| R $\alpha$ RagC | IF | 1:400 | Cell Signaling | 3360 |
| **Secondary Antibody** | **Use** | **Dilution** | **Company** | **Catalog Number** |
| IRDye® 800CW G $\alpha$ M - IgG | WB | 1:5000 | LI-COR | 925-32210 |
| IRDye® 800CW G $\alpha$ R - IgG | WB | 1:5000 | LI-COR | 925-32211 |
| IRDye® 680CW G $\alpha$ R - IgG | WB | 1:5000 | LI-COR | 925-68071 |
| IRDye® 680CW D $\alpha$ G - IgG | WB | 1:5000 | Invitrogen | A-21084 |
| D $\alpha$ G-ALEXA488 | IF | 1:250 | ThermoFisher | A-11055 |
| D $\alpha$ R-ALEXA488 | IF | 1:250 | ThermoFisher | A-21206 |
| R $\alpha$ M-ALEXA488 | IF | 1:250 | ThermoFisher | A-11061 |
| G $\alpha$ M-ALEXA488 | IF | 1:250 | ThermoFisher | A-11029 |
| D = Donkey, G = Goat, M = Mouse, R = Rabbit |  |  |  |  |
| **Reagent** | **Use** | **Dilution** | **Company** | **Catalog Number** |
| Prolong Diamond w/ DAPI | Antifade/IF |  | Thermofisher | P36971 |
| Torin-1 | mTOR inhibitor | 250nM | Selleckchem | S2827 |
| Vacuolin-1 | Increase endosomal size | 5μM | Sigma-Aldrich | 673000 |

### Cell culture and light microscopy

All cells used in this study were cultured in High Glucose Dulbecco’s Modified Eagle’s Medium (Invitrogen) supplemented with 10% FCS and 1% penicillin/streptomycin (Invitrogen) in a 5% CO2-humidified incubator at 37°C. To induce autophagy, cells were starved for 2 hours at 37°C using EBSS (ThermoFisher) and, where indicated, restimulated for 15 min with complete medium. Cells were transiently transfected using X-tremeGENE™ (Roche) according to manufacturer’s instructions. The generation of the CRISPR/Cas9 knockout cells is described in(Van der Welle et al., 2021). Generation of stable cell lines was achieved through a minitol-transposase system. The desired genes (e.g. VPS41-GFP) were cloned into donor vectors with puromycin resistance under an EF1α promotor. These donor constructs were coexpressed with a transposase in 1:2 ratio for genomic integration using Effectene transfection reagent (301425, QIAGEN). After 24 hours, the cells were put on puromycin selection for 7-14 days to ensure stable genomic integration.

For immunofluorescence microscopy, cells were washed with PBS once and fixed using 4% w/v paraformaldehyde (PFA, Polysciences Inc.) for 20 min. Cells were washed 3 times with PBS, permeabilized with 0.1% Triton X-100 (Sigma) for 5 min and blocked for 15 min using a 1% BSA solution. Samples were labeled, embedded in Prolong DAPI (Invitrogen) and imaged on a Leica Thunder fluorescence microscope using a 100x, 1.47 NA oil objective, a Photometrics Prime 95B scMOS camera and LAS X software. For transfected samples, cells with similar transfection levels were selected. For TFE3 localization in Fig 1C, D and Fig EV1A, cells were scored positive when a defined nuclear outline was visible due to nuclear labeling of TFE3. RagC-LAMP1 co-localization was quantified using object detection-based analysis in the ComDet plugin for Fiji (Eugene Katrukha, Cell biology, Utrecht University) and a custom macro.

### Co-immunoprecipitation and Western Blotting

Cells were seeded in 15cm dishes and, when appropriate, transfected with the constructs as indicated in the respective figures and according to above mentioned transfection protocols. Cells were washed 3 times with ice-cold PBS and lysed using a CHAPS lysis buffer (50mM Tris pH=7.5, 150mM NaCl, 5mM MgCl2, 1mM DTT, 1% (w/v) CHAPS) complemented with protease and phosphatase inhibitor (Roche). Cells were collected by scraping and then spun down at 13000rpm for 15 min at 4°C. Protein levels were equalized between samples using a Bradford Protein Assay (Bio-Rad). 10% of the sample was saved as input control. The remaining lysate was incubated for 1 hour with uncoated protein G beads (Millipore) to remove a-specifically bound protein. Beads were spun down and the supernatant was incubated with beads together with 2µg antibody against the prey. GFP-Trap beads (Chromotek) were used for immunoprecipitation of GFP-tagged constructs. Samples were incubated overnight, extensively washed and eluted using SDS sample buffer and run on precast gradients (4-15%), or 10% gels (Bio-Rad). Gels were transferred using Trans-Blot® Turbo™ RTA Mini PVDF Transfer Kit and the Trans-Blot® Turbo™ Transfer system (Bio-Rad). Membranes were blocked with Odyssey® Blocking Buffer (LI-COR) in 0.1% TBST for 1 hour at RT and incubated with primary antibody diluted in Odyssey® Blocking Buffer (LI-COR) in 0.1% TBST overnight at 4°C, rocking. Membranes were washed extensively with 0.1% TBST and incubated with secondary antibodies, diluted in Odyssey® Blocking Buffer (LI-COR) in 0.1% TBST at RT for 1 hour, rocking. The membranes were again washed extensively with 0.1% TBST, followed by 2 washing steps with PBS. Membranes were scanned using the Amersham™ Typhoon™ Laser Scanner (GE Healthcare Life Sciences).

### Plasmids

The generation of the GFP-VPS18, VPS39-V5, VPS41-GFP, VPS41^285P^-GFP and VPS41^R662*^-GFP constructs is described in (Van der Welle et al., 2021). The constructs encoding for HA-GST-RagA^21L^/RagA^66L^, HA-GST-RagB^54L^/RagB^99L^, HA-GST-RagC^75L^/RagC^120L^, HA-GST-RagD^77L^/RagD^121L^ were kindly provided by Dr. F.J. Zwartkruis, University Medical Center Utrecht). FLAG-RHEB^N153T^ and FLAG-FLCN were obtained from Addgene (plasmid #19997 and #72290 respectively) (Urano et al., 2007). Minitol vectors were generated from the previously mentioned VPS41-GFP constructs and a pMinitol2-hEF1a-MCS-polyA-pgk-Puro backbone using XhoI and NotI.

### Statistics

Quantification of all western blots was performed using Fiji software (Schindelin et al., 2012). Two-tailed Students t tests were performed to compare two samples whereas ANOVA was performed for comparison of multiple samples to analyze statistical significance. Data distribution was assumed to be normal, but this was not formally tested. All error bars represent the Standard Error of the Mean (SEM) as indicated in the specific figure legends.

## Data availability

This study includes no data deposited in external repositories.

## Appendix

We thank our colleagues of the Cell Biology section and Center for Molecular Medicine for the fruitful discussions and feedback. We especially thank Prof. A. Ballabio and Dr. G. Napolitano for their insightful comments and valuable input.

NL is supported by a ZonMW TOP grant [40-00812-98-16006 to J.K]. RW and PS are supported by a DFG grant [FOR2625 to J.K. as part of the Research Consortium]

Author contribution: Experiments: RW, JB, PS; Study design: RW, JB, PS, NL and JK; Manuscript reviewing and revision: FZ, JB, PS, NL; Study design and supervision: RW and JK; Funding: JK; Manuscript writing: RW and JK.

The authors declare no conflict of interest.

