## Supplemental figures for "HOPS is required for non-canonical mTORC1 signaling by recruiting Rags and FLCN to lysosomal membranes"

Supplemental Figure 1

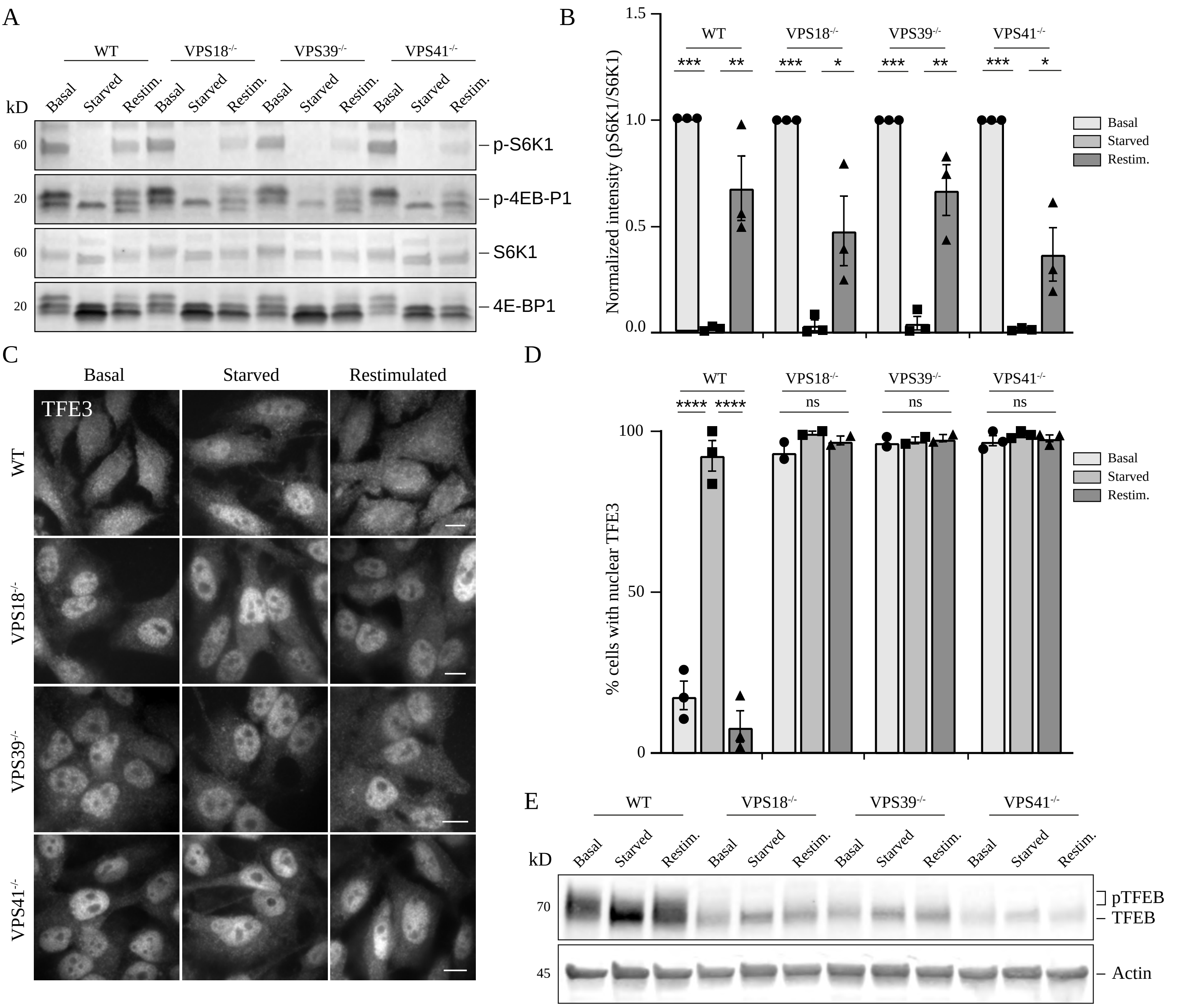

**Supplemental Figure 1. Depletion of HOPS subunits results in impaired phosphorylation of TFE3 and TFEB**

A. Western blot showing the phosphorylation status of mTORC1 substrates S6K1 and 4E-BP1 under basal (untreated, fed), starved (2 hours) or restimulated (15 min) conditions. Phosphorylation in VPS18$^{-/-}$, VPS39$^{-/-}$ or VPS41$^{-/-}$ cells was similar as in WT cells.

B. Quantification of A, numbers represent mean ± SEM, *P < 0.05, **P < 0.01, ***P < $10^{-4}$. One-way ANOVA with Bonferroni correction) (n=3).

C. Immunofluorescence of endogenous TFE3 in WT, VPS18$^{-/-}$, VPS39$^{-/-}$ or VPS41$^{-/-}$cells under basal (untreated, fed), starved (2 hours) and restimulated (15 min) conditions (Scale bar, 10 µm, n=3).

D. Percentage of cells with nuclear TFE3 localization (mean ± SEM, ****P < $10^{-5}$. One-way ANOVA with Bonferroni correction). In contrast to WT cells, HOPS depleted cells showed continuous nuclear localization of TFE3.

E. Western blot showing the phosphorylation status of TFEB in WT, VPS18$^{-/-}$, VPS39$^{-/-}$ and VPS41$^{-/-}$ cells under basal (untreated, fed), starved (2 hours) or restimulated (15 min) conditions. Phosphorylation of TFEB is visualized by a smear, whereas a distinct band indicates dephosphorylated TFEB, reflecting nuclear translocation. WT cells showed a smear under basal and restimulated conditions and a distinct band upon starvation. VPS18$^{-/-}$, VPS39$^{-/-}$ and VPS41$^{-/-}$ cells showed a distinct band regardless of nutrient availability. (n=3).

Supplemental Figure 2

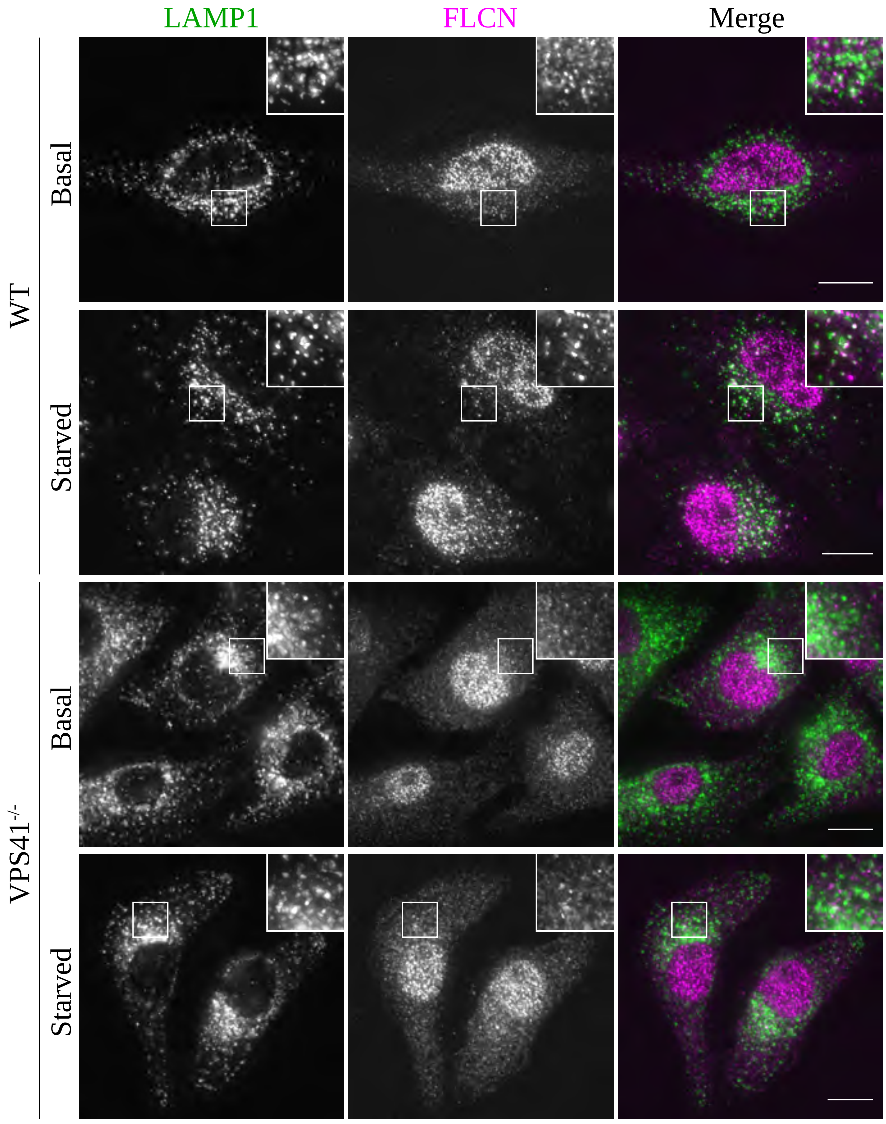

**Supplemental Figure 2. Lysosomal localization of FLCN is impaired in VPS41$^{-/-}$ cells**

Immunofluorescence of endogenous FLCN and LAMP1 in WT and VPS41$^{-/-}$ cells under basal and starved (2 hours) conditions. In WT cells, FLCN and LAMP1 co-localized upon starvation. In VPS41$^{-/-}$ cells, no localization was observed. (Scale bar, 10 µm, insert, 8 µm). (n=3).

Supplemental Figure 3

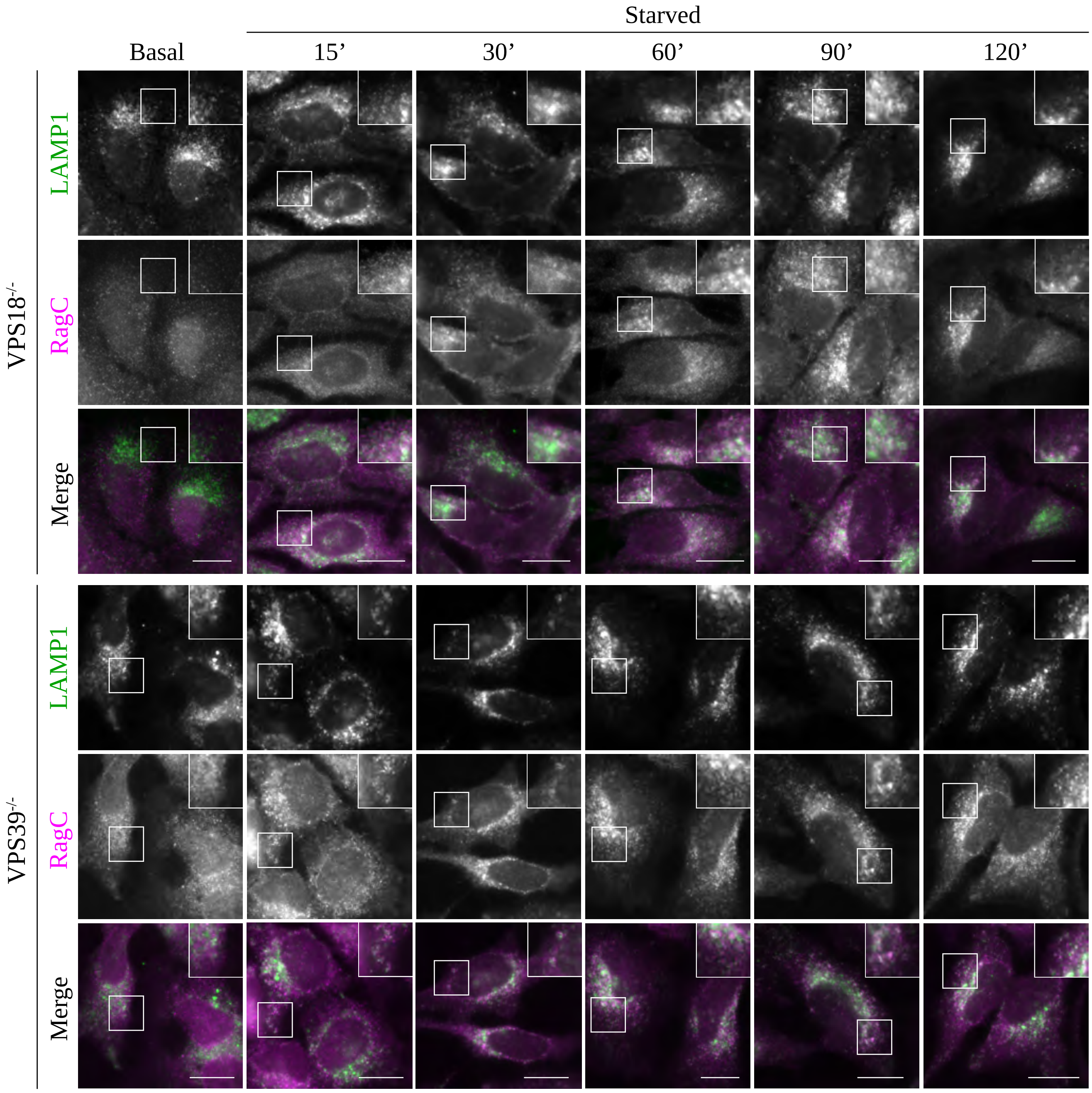

**Supplemental Figure 3. The HOPS complex is required for lysosomal recruitment of RagC**

Immunofluorescence of endogenous RagC and LAMP1 in VPS18$^{-/-}$ and VPS39$^{-/-}$ cells under basal conditions and after different time points of starvation. In VPS18$^{-/-}$ and VPS39$^{-/-}$ cells, RagC remained cytosolic, showing little colocalization with LAMP1. Similar results were obtained for VPS41$^{-/-}$ (Fig 3). (Scale bar and inserts, 15µm, n=2).

Supplemental Figure 4

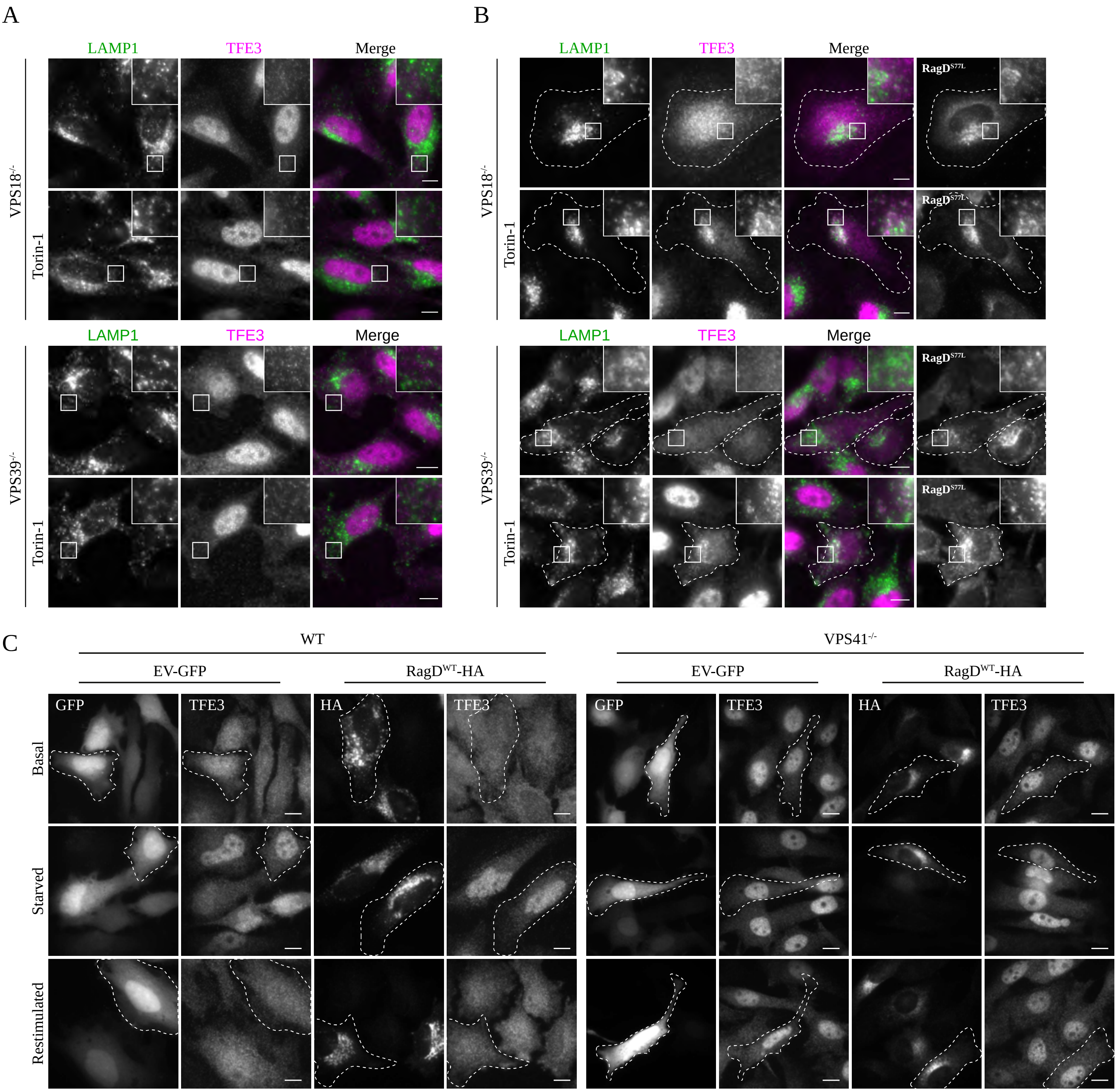

**Supplemental Figure 4. Active RagD rescues the TFE3 phenotype in VPS18$^{-/-}$ and VPS39$^{-/-}$ cells.**

A, B. Immunofluorescence of endogenous TFE3 and LAMP1 in VPS18$^{-/-}$ or VPS39$^{-/-}$ cells in the presence or absence of 250nM Torin-1 and upon transient transfection of constitutively active RagD (RagD$^{S77L}$). Overexpression of RagD$^{S77L}$ in VPS18$^{-/-}$ and VPS39$^{-/-}$ cells resulted in cytosolic retention of TFE3, and upon Torin-1 treatment increased co-localization with LAMP1 and RagD. Similar results were obtained for VPS41$^{-/-}$ cells (Fig 4B) (transfected cells visualized by white outline). (Scale bar and insert, 10µm. n=3).

C. Immunofluorescence of endogenous TFE3 in WT or VPS41$^{-/-}$ cells under basal, starved (2 hours) or restimulated (15 min) conditions upon expression of EV-GFP or RagD$^{WT}$-HA. WT cells show TFE3 in the nucleus only upon starvation, regardless of RagD$^{WT}$ expression. TFE3 is constitutively localization to the nucleus in VPS41$^{-/-}$ cells. Expression of RagD$^{WT}$ did not rescue this phenotype.

Supplemental Figure 5

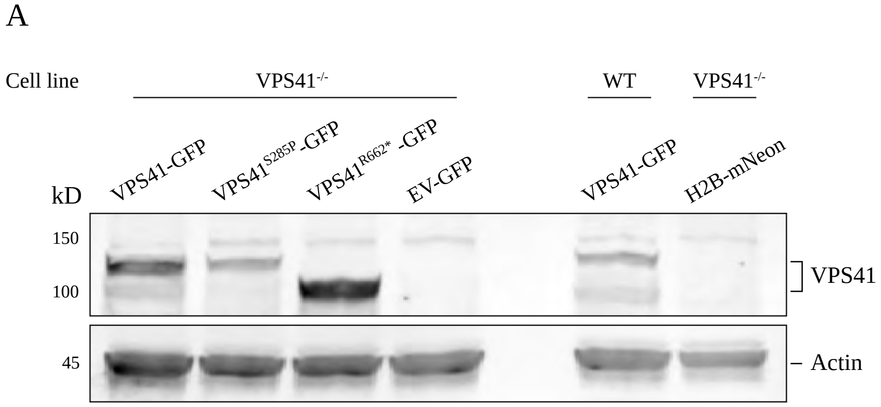


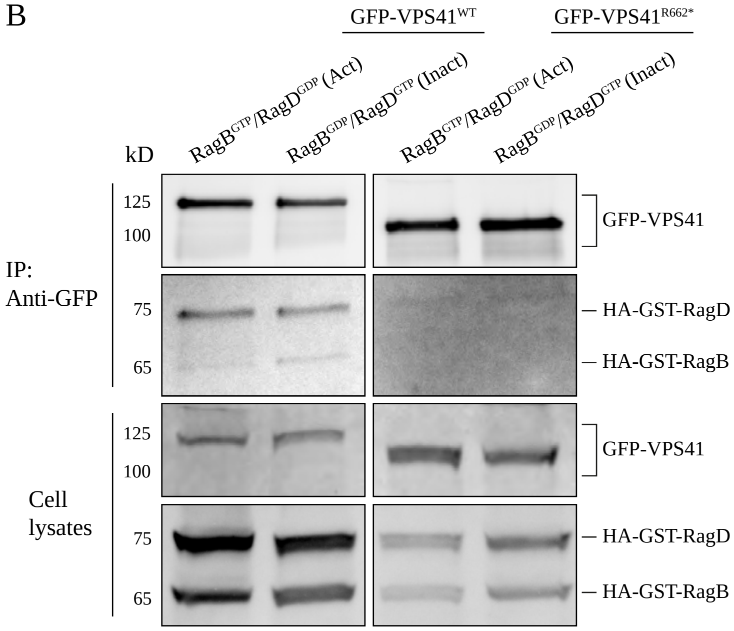

**Supplemental Figure 5.**

A. Western Blot showing the stable expression of $VPS41^{WT}$-GFP, $VPS41^{S285P}$-GFP and $VPS41^{R662*}$-GFP in $VPS41^{-/-}$ cells as well as the expression of $VPS41^{WT}$-GFP in WT cells. H2B-mNeon serves as a negative control.

B. Co-immunoprecipitation of cells stably expressing $VPS41^{WT}$-GFP or $VPS41^{R662*}$-GFP and transiently transfected with either $RagB^{GTP}$/$RagD^{GDP}$ (active conformation) or $RagB^{GDP}$/$RagD^{GTP}$ (inactive conformation). $VPS41^{WT}$ interacts with both the active and inactive Rag heterodimers. The $VPS41^{R662*}$ mutant, which lacks the RING domain, showed severely reduced affinity for the RagB/RagD heterodimer, regardless of conformation (n=3).
